# Recombinant Expression of Metal-Binding Proteins (Bacterioferritin, MBP, Metallothionein, and OprF) and Pesticide-Binding Proteins (LC-Cutinase and OpdA) in Escherichia coli for Detection of Agricultural Contaminants

**DOI:** 10.1101/2020.06.04.135665

**Authors:** Kimberly Hwang, Allison Kuo, Eugene Choi, Jessica Chen, Jonathan Hsu, Jude Clapper

**Author notes:** Financial Disclosure This work was funded by Taipei American School. The funders had no role in study design, data collection, and analysis, decision to publish, or preparation of the manuscript.

## Abstract

We consume fruits and vegetables every day without knowing whether or not agricultural residues (i.e. pesticides & heavy metals) are present on them. Current methods of agricultural residue detection are not easily accessible to the public and are inconvenient for everyday use. Thus, we want to develop a convenient visualization of agricultural residues by designing metal-binding and pesticide-binding proteins along with colored proteins that can directly interact with these residues. We worked with metal-binding proteins including Bacterioferritin, MBP, Metallothionein, and OprF. We also utilized pesticide-binding proteins LC Cutinase and OpdA. In this paper, we envision a system in which our residue-binding proteins are fused with chromoproteins to allow for the visible detection of agricultural residues on produce. Our final product can be used by consumers, distributors and farmers alike.

## Introduction

### The Problem With Agricultural Contaminants

Agricultural contaminants, such as heavy metals and pesticides, are chemical substances remaining in crops that are harmful to human health (Agriculture Victoria, n.d.). Heavy metals, such as copper, mercury, nickel, iron, lead, and cadmium, are commonly found in industrial waste runoff that contaminates farmland (Yang et al., 2018; Zhang et al., 2013; Jaishankar et al., 2014; Temple & Bisessar, 1981). Pesticides, such as insecticides, herbicides, fungicides, are substances that kill, repel, or control pests to maximize crop yields (Pesticide Action Network, 2015).

In general, exposure to heavy metals has detrimental health effects, including high blood pressure, renal dysfunction, brain damage, or death (Jaishankar et al., 2014). In 2016, lead exposure was responsible for approximately 540,000 deaths worldwide (World Health Organization, 2018).

Moreover, chronic exposure to pesticides can lead to cancer, Alzheimer’s disease, Parkinson’s disease, endocrine disorders, development disorders, infertility, and even death (World Health Organization, 2018). In 2017, exposure to pesticides caused 200,000-300,000 deaths worldwide (UN News, 2017).

### Current Detection Methods

Given the prevalence of heavy metals and pesticides in our home country of Taiwan and all over the world, we developed a simple and convenient method to detect agricultural contaminants on food products.

Current lab methods to detect heavy metals, such as spectrometry, are usually expensive, complicated, and time-consuming (Sikdar, 2018). Though there are home testing kits available, these kits vary in accuracy and most lack approval from official governmental bodies. For pesticide residue detection, farmers typically send samples to pesticide-detection companies, such as Société Générale de Surveillance (SGS), that will help test residue levels on the samples using gas or liquid chromatography (Société Générale de Surveillance, 2019). However, our research indicated that most farmers are unaware of the exact details behind lab testing, and this process does not provide immediate results for farmers.

Our product involves a liquid spray containing colored residue-binding proteins; when applied to food products, the proteins will bind to agricultural contaminants and produce patches of color that will remain on the contaminated regions (Figure 4).

**Figure 1.**
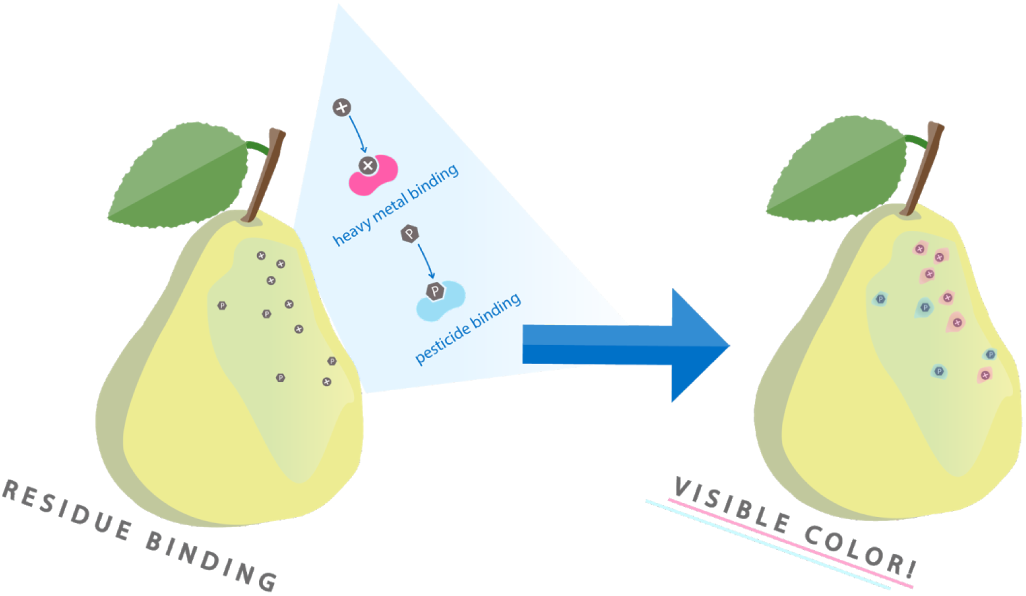
Protein Binding to Residues. We envision that the colored residue-binding proteins will be applied to food products and hence visualize the contaminated regions.

## Materials/Methods

### Metal-Binding Protein Constructs

Our metal-binding proteins include bacterioferritin, Metal Binding Protein (MBP), Metallothionein, and OprF. Bacterioferritin is a natural bacteria iron storage protein and MBP is a lead-binding protein derived from PbrR expressed in the cytosol. Metallothionein can tightly chelate metal ions by forming a strong coordination bond and OprF+CBP is a nonspecific membrane porin for small hydrophilic molecules fused with a copper-binding peptide.

### Pesticide-Binding Protein Constructs

Our construct of pesticide-binding proteins consisted of LC Cutinase, a fungal enzyme that degrades cutin, water-soluble esters and insoluble triglycerides, and OpdA, a phosphotriesterase that can detoxify a broad range of organophosphate pesticides.

### DNA Design and Protein Expression

**Figure 2.**
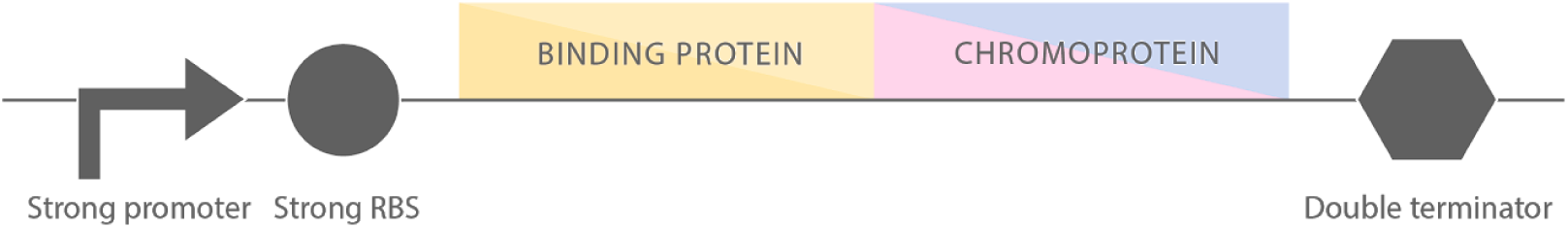
Construct design of colored residue-binding protein. For each of our binding proteins, we inserted an upstream constitutive strong promoter-RBS combination and a downstream color binding protein, followed by a strong double terminator to the construct to maximize protein expression.

To ensure that the binding protein and colored protein were fused, we inserted either a flexible glycine-serine linker (GS) or a more rigid EAAAK linker between the binding protein and the chromoprotein. For protein purification in downstream applications, we included a hexahistidine tag (6xHis) after the RBS and before the ORF. These constructs were synthesized by Twist Bioscience (henceforth called Twist constructs). PCR check and Tri-I Biotech sequencing results indicated that we had successfully cloned these constructs.

### Metal-Binding Assay

Our metal-binding parts produce intermembrane proteins expected to increase the cells’ capacity to store heavy metal ions. To test the functionality of our proteins, we detected the difference in the ion storage capacity of construct-expressing cells and negative-control cells. Thus, we had two experimental groups: cells expressing the fusion protein and cells expressing the binding protein only. We had two negative control groups: cells expressing RFP only and cells carrying an ORF with no promoter (ORF only) plasmid.

In order to measure cell storage capacity, we incubated cells with the targeted heavy metal ions and then measured the absorbance of extracellular metal ions. By the Beer-Lambert law, concentration is directly proportional to absorbance. For the experimental groups, we expected the extracellular solution to have a lower concentration of metal ions and, thus, a lower absorbance as compared to the negative control.

Metal solutions were prepared by dissolving the appropriate metal ions into distilled water. To optimize the absorbance measurements in the downstream experiment, the wavelength at the peak absorbance of metal solutions were first determined using a spectrophotometer.

We prepared and standardized overnight bacterial cultures to an OD600 of 0.7. Then, we centrifuged the cultures and resuspended the pellet in heavy metal solutions. We gently shook the cell-heavy metal mixtures at room temperature for approximately 2 hours. We then spun down the cells to isolate the extracellular solution as the supernatant. We measured the peak absorbance of the supernatant using a spectrophotometer.

**Figure 3.**
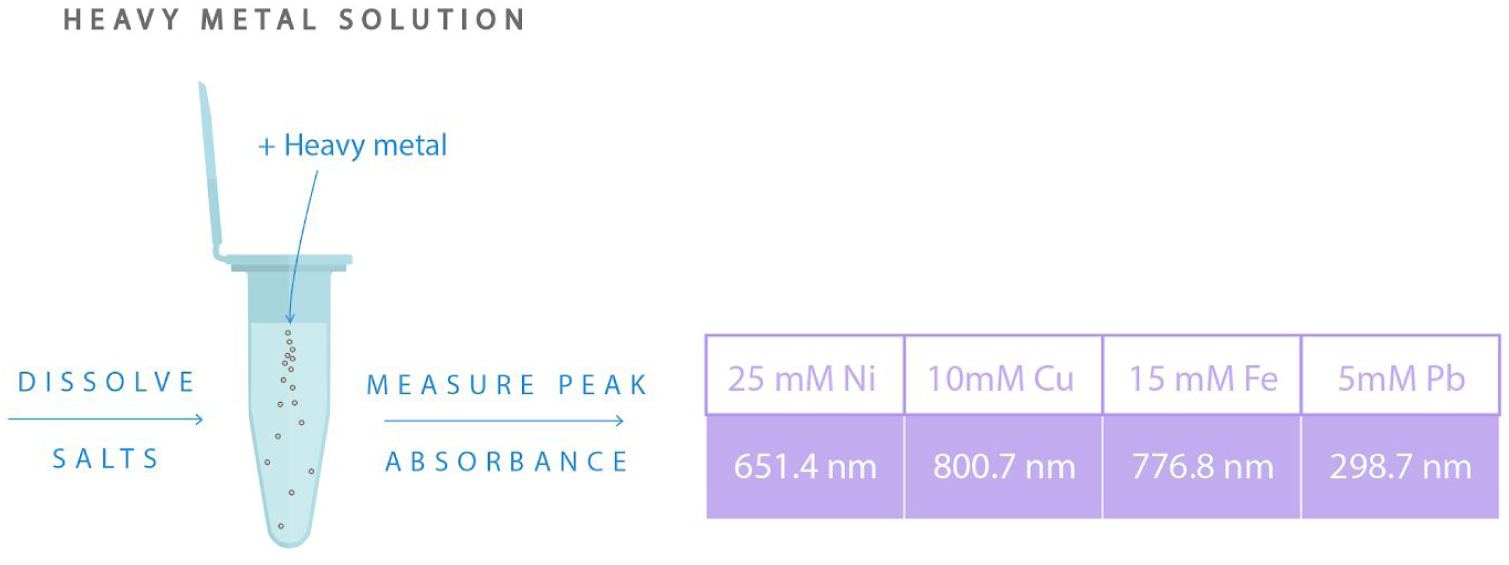
The wavelength for peak absorbance of heavy metal ion solutions. First we prepared bacterial cultures to an OD600 of 0.7. We centrifuged the cultures and resuspended the pellet in various heavy metal solutions. Then, we shook the cell-metal mixture at room temperature for around 2 hours and centrifuged the mixture. We measured the peak absorbance of the supernatant which contained the extracellular solution using a spectrophotometer.

### Pesticide Binding Enzyme Assay: pH Test

Our pesticide-binding construct, LC-Cutinase, produces intermembrane enzymes expected to break malathion into malathion monoacid and malathion diacid. This means that a functional enzyme would produce an acidic degradation product and thus lower the pH of the solution. To prove that malathion degradation decreases pH, we hydrolyzed Malathion under a high pH environment.

**Figure 4.**
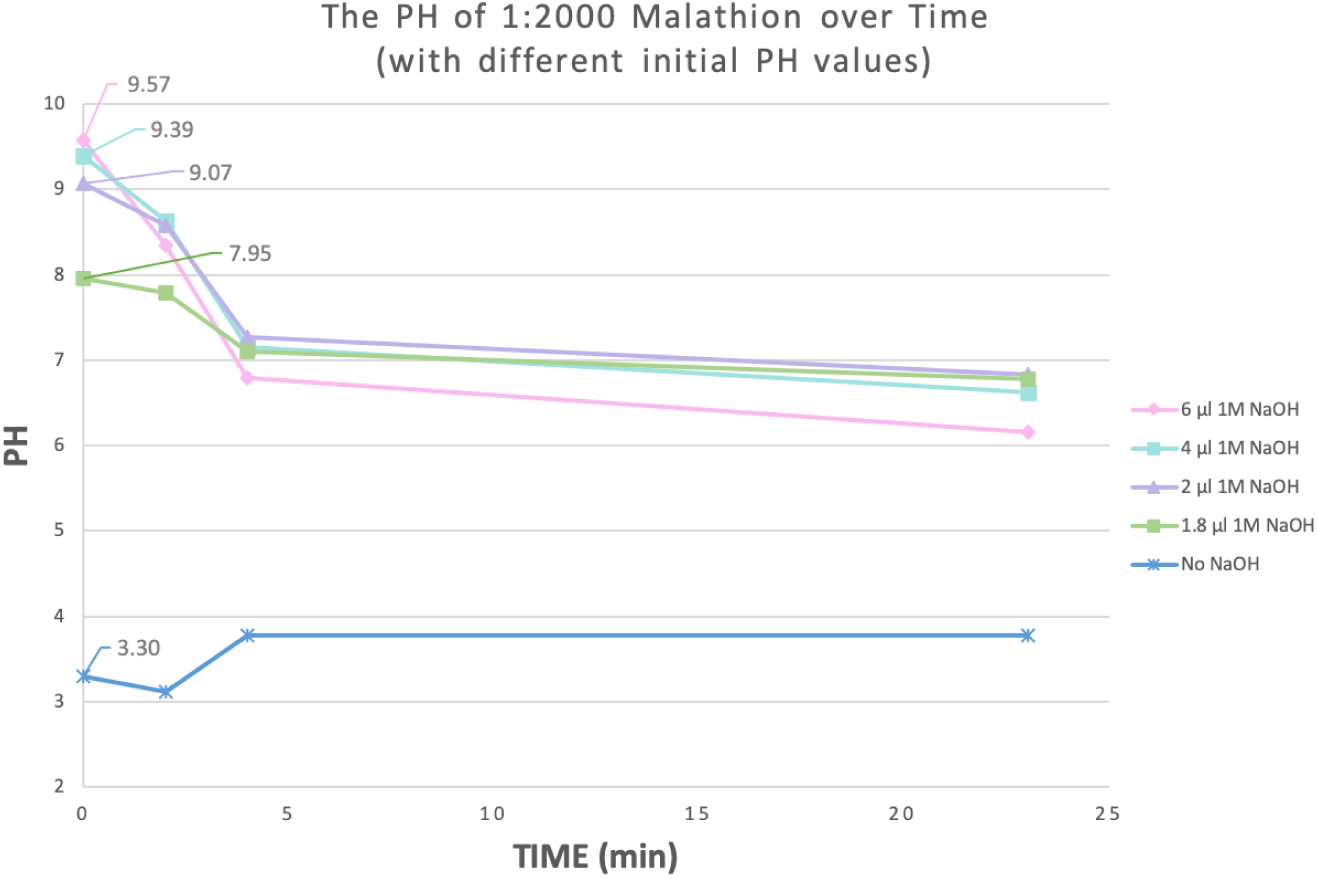
Malathion’s natural degradation in H2O decreases pH. After adding NaOH to increase the pH malathion to different levels, we measured the pH of malathion over time. We observed that malathion naturally decreases in pH only when it has a high, basic initial pH.

Our results show that, under high pH, malathion degrades and lowers in pH. Thus, malathion does break down into the two acids to release protons and lower the pH. To test the functionality of our proteins, we compared the difference in pH levels of solutions of construct-expressing cells and negative-control cells. Thus, we had an experimental set: cells expressing the LC-Cutinase-GS-AmilCP protein, and two negative control groups: cells expressing amilCP only and cells carrying an ORF-only plasmid.

Overnight bacterial cultures were prepared and standardized to an OD of around 0.7. Then, the cultures were centrifuged and the pellet was resuspended in 2 mL of 1:1000 diluted malathion solution. The pH of each solution was measured every 5 minutes over 24 hours.

### Pesticide Binding Protein Assay: Ellman’s Test

The protein OpdA cleaves malathion to expose its sulfhydryl group (Scott et al., 1970). We used Ellman’s reagent, which reacts with thiol groups like sulfhydryl to produce a yellow color (ThermoFisher, 2011). The amount of sulfhydryl can be quantified by measuring the absorbance at 412.4 nm. We decided to use Ellman’s to determine the functionality of our OpdA constructs. Additionally, the change in color would mimic the original idea of a visual detection of pesticides.

Through literature research, we found that Ellman’s reagent reacts on a 1:1 molar ratio with malathion. Using the molar weight of both compounds, we calculated the concentration and volume ratio needed for Ellman’s. As a preliminary test, we prepared cell lysate from cells constitutively expressing our OpdA construct. The OpdA cell lysate was then added to either H2O or malathion. Our results showed that only when malathion was added did we see a noticeable increase in yellow color, both by eye and by absorbance at 412 nm.

**Figure 5.**
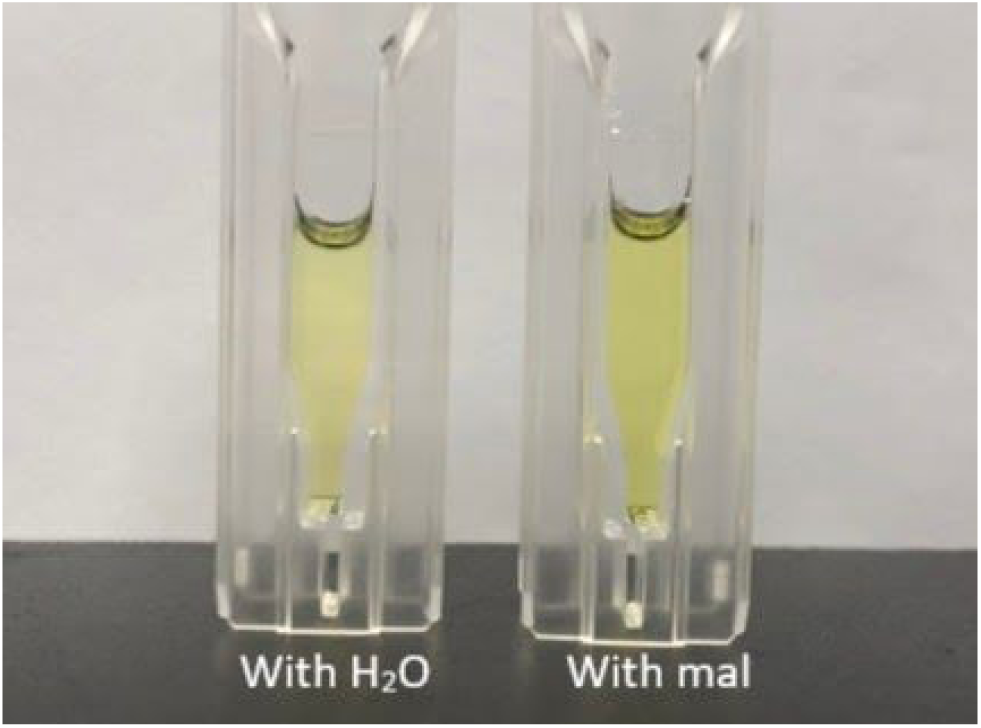
Visual comparison of OpdA + mal and OpdA + H2O only with Ellman’s added. This comparison shows a distinct color difference between a OpdA + mal sample and OpdA + H2O with Ellman’s reagent sample, indicating the proper functioning of the constructs

**Figure 6.**
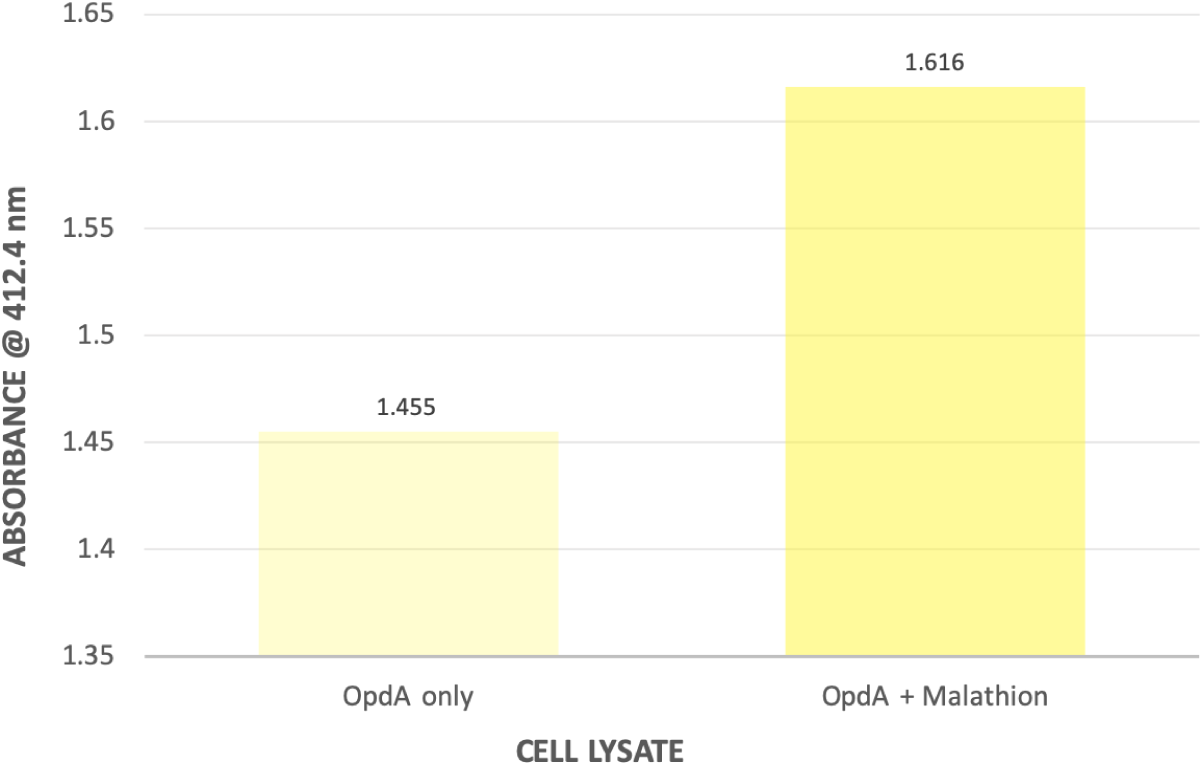
Absorbance at 412.4nm for OpdA + mal and OpdA + H2O, with Ellman’s added. OpdA + mal has a higher absorbance value, which shows that OpdA cleaves malathion to expose thiol groups, which react with Ellman’s to turn more yellow.

Addition of malathion with purified OpdA resulted in a darker yellow compared to when water was added. Utilizing the 6xHis tag on our OpdA construct, we purified the proteins using the His GraviTrap kit (GE Healthcare). We confirmed the presence of our purified proteins by SDS-PAGE.

**Figure 7.**
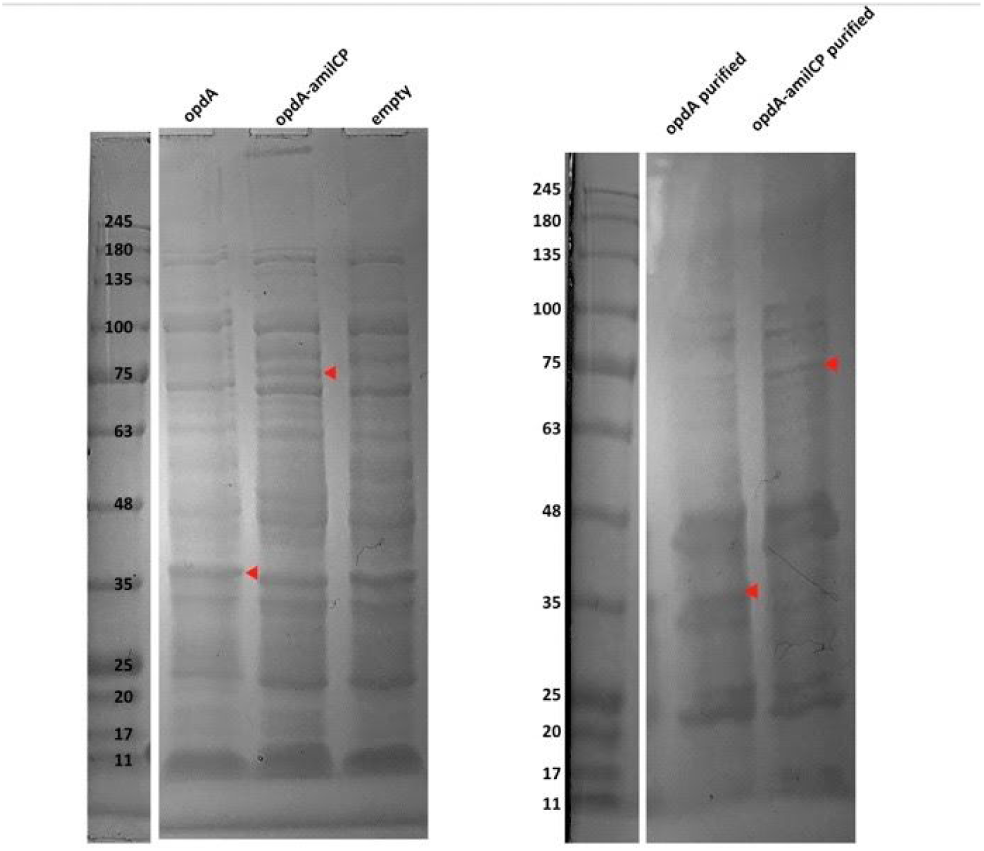
Protein gel of purified OpdA and OpdA-amilCP proteins. To verify OpdA expression in E. coli, we subjected OpdA lysate (left gel) and OpdA purified protein (right gel) to SDS-PAGE, expecting a signal at around 37 kDa. On the left gel, we saw a strong signal at around 37 kDa in the OpdA lane, but not in the empty lane that was used as a control. On the right gel, we also saw a strong signal at around 37 kDa in the OpdA purified lane.

We used malathion + elution buffer (EB) from Geneaid as our control since our experimental sets also have malathion and EB but with the addition of proteins, produced by our constructs. In order to mitigate the color of our control, we needed to see if the proteins have an additional effect on the yellow color. We found that the more diluted the protein concentration, the fainter the yellow, which meant our purified proteins contributed more to the yellow color. This also gave us a confirmation that OpdA functions as expected.

**Figure 8.**
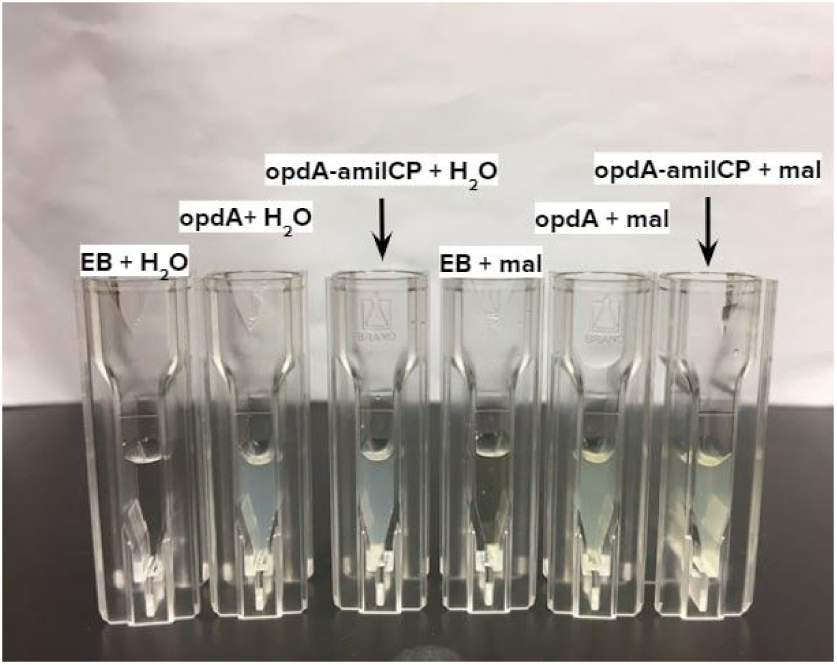
Visual test with Ellman’s Reagent added. Every cuvette has the volume ratio of 225uL EB/OpdA/OpdA-amilCP + 150uL H2O/mal + 150uL Ellman’s reagent. The two on the right look the most yellow, which is expected, as they have our pesticide-binding proteins and malathion.

To fully test the functionality of OpdA, we ran a time trial of the reaction between OpdA and malathion. We expected that over time, the yellow absorbance would go up, as OpdA cleaves more malathion. We took absorbance data at 412nm at 2-minute intervals for a total of 10 minutes, and at 20 and 30 minutes. Initially, we found that the experimental set turned yellow immediately, and its absorbance values either did not increase or even decreased over time, which suggested the enzymes were so efficient in cleaving the malathion that is was difficult for us to detect an increase in absorbance to model enzyme kinetics. This is great for our final product because we were interested in an immediate color detection of contaminants, but it was hard for us to model the enzyme kinetics. Thus, we diluted our original purified proteins by 1:1000 and repeated the procedure.

## Results and Discussion

**Figure 9.**
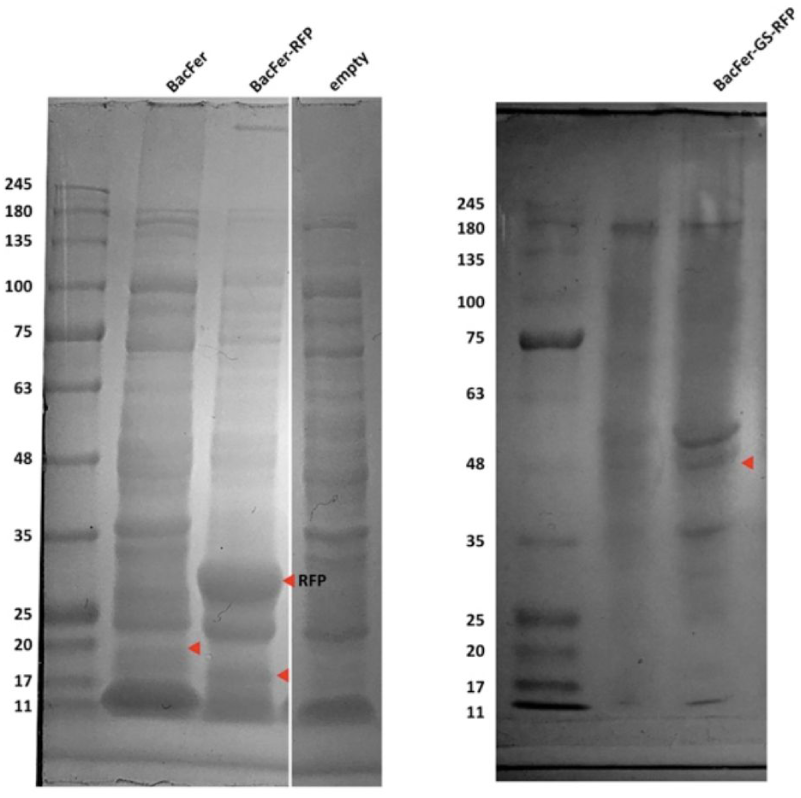
Protein Gel of BacFerr-mRFP. To verify BacFerr-mRFP expression in E. coli, we subjected BacFerr-mRFP lysate to SDS-PAGE, expecting a signal at around 45 kDa. Instead, we saw separate signals at approximately 18 kDa and 26 kDa in the BacFerr-mRFP lane. We think that the BacFerr migrated at a smaller size compared to BacFerr only due to partial proteolytic cleavage. Seeing that the BacFerr and mRFP were expressed separately, we designed a DNA construct (BBa_K2921350) in which we added a GS-linker to connect the two proteins.

The results of our Coomassie-stained protein gels of our constructs indicated the presence of the binding proteins and colored proteins, suggesting that the proteins were expressed separately, rather than fused together (as seen in the left gel pic in Figure 9). Since the binding proteins themselves were still expressed, these constructs were still used for preliminary assays.

The results of our Coomassie-stained protein gels of our constructs with the addition of GS or EAAAK linkers indicated the presence of the binding proteins and colored proteins (as seen in the right gel pic in Figure 9). The protein signals suggested that the proteins were expressed as a single, fused unit. Additionally, the lack of a protein signal at the chromoprotein band further suggested that the proteins were not separate, indicating that the linker sequence successfully fused the binding protein to the chromoprotein.

### Metal-Binding Assay

Our results indicate that there are lower concentrations of metal ions in the extracellular supernatants of cells expressing the fusion protein and the binding protein, as compared to the negative controls. This shows that our fusion and binding proteins are capable of binding to their targeted metal ions, thus increasing the cell’s ability to retain metal ions from their environment.

**Figure 8.**
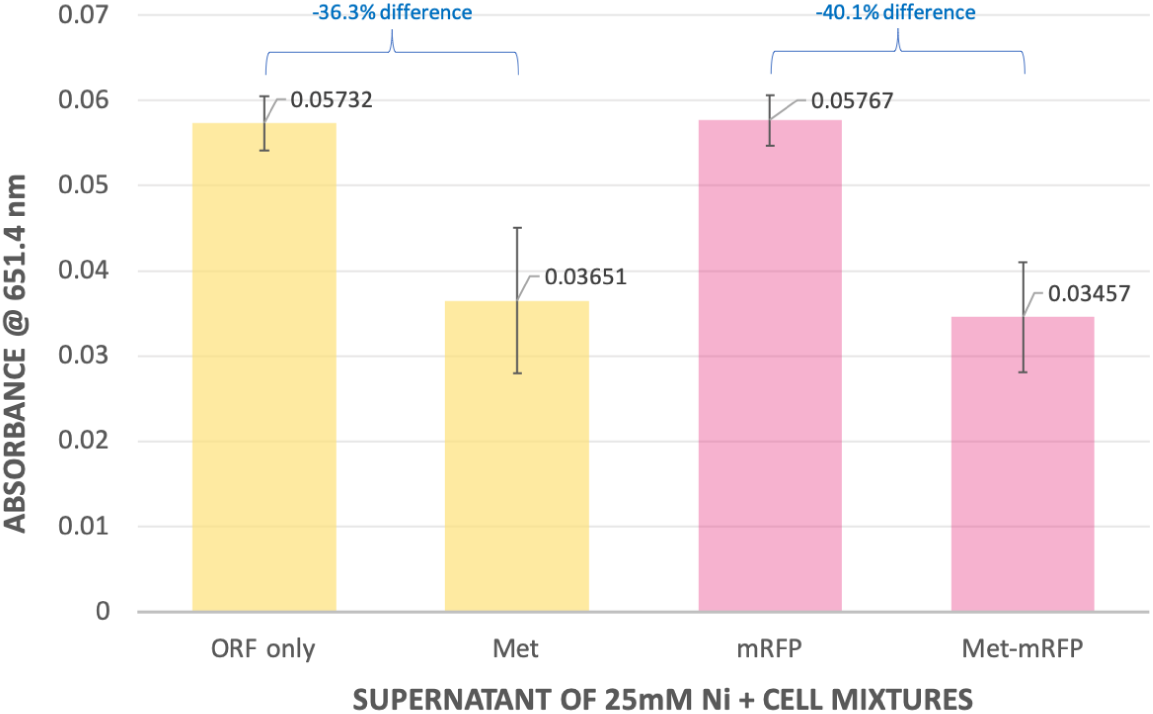
Metallothionein and Metallothionein-RFP increase cellular retention of nickel ions. The graph shows that our binding proteins are capable of binding to their targeted metal ions and increasing the cell’s ability to retain metal ions.

After two hours of shaking incubation with nickel ions, we centrifuged all samples to isolate the extracellular solution. At 651.4 nm, we observed lower absorbance in extracellular solution of cells expressing Metallothionein and Metallothionein-RFP compared to the negative control cells. We repeated the same metal-binding test with fusion proteins synthesized by Twist Bioscience, which included a linker sequence to fuse the binding protein to mRFP. We used cells expressing mRFP only as the negative control. We found that for all of our Twist constructs except MBP, there are lower concentrations of metal ions in the extracellular supernatants of cells expressing the fused proteins, as compared to negative control. These results suggested that our fused proteins are capable of binding to their targeted metal ions, thus increasing the cell’s ability to retain metal ions from their environment.

**Figure 9.**
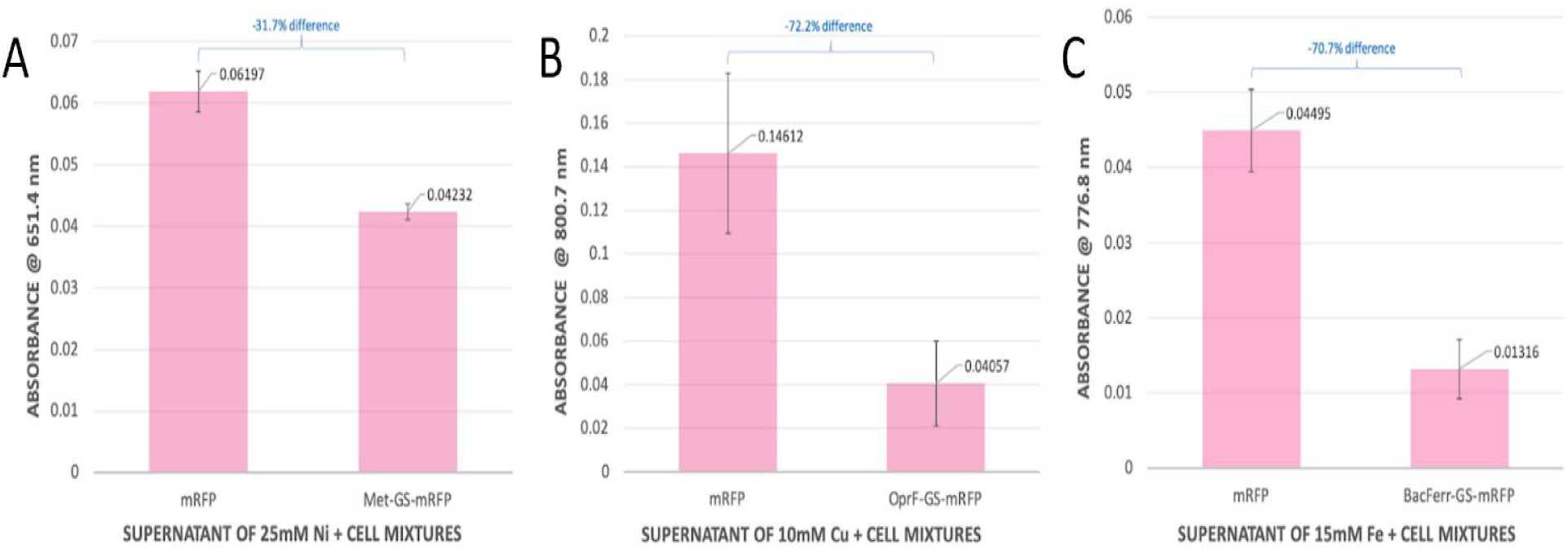
Twist Metal-Binding Fusion Proteins (with GS-linker) increases cellular retention of heavy metal ions. **A)** We observed a lower absorbance in the extracellular solution of cells expressing the fusion proteins compared to the negative control cells for the MET-GS-mRFP construct. **B)** We observed a lower absorbance in the extracellular solution of cells expressing the fusion proteins compared to the negative control cells for the OprF-GS-mRFP construct. **C)** We observed a lower absorbance in the extracellular solution of cells expressing the fusion proteins compared to the negative control cells for the BacFerr-GS-mRFP construct.

After two hours of shaking incubation with nickel ions, we centrifuged all samples to isolate the extracellular solution. We observed lower absorbance in the extracellular solution of cells expressing the fusion proteins compared to the negative control cells. We did not test our twist MBP-GS-mRFP construct because during our preliminary metal-binding test with our cloned MBP and MBP-mRFP constructs, we observed immediate clumping of the cell-lead mixtures, which resulted in discrepancies in our results. The clumping is most likely a sign of bacteria death in the presence of a high concentration of lead.

## Conclusion

Preliminary analysis of the time-based data measuring the pH of LC cutinase at room temperature over time suggested that LC cutinase was either not functioning correctly due to being very inefficient or not present. Through more research, we discovered that the optimum condition for LC cutinase is at a pH of 8.5 and a temperature of 50°C (Sulaiman, et al., 2011). We tried these experiments at 50°C and even at optimal conditions, we did not see an improvement in function. This led us to suspect that the enzyme was not actually present in the experiment. When we checked for the presence of our protein by SDS-PAGE, we did not see bands at the correct size, suggesting that LC-Cutinase might not have been expressed in the cells.

Utilizing *Escherichia coli* (*E. coli*) as our chassis, we synthesized a series of chromoprotein-fused proteins designed to bind to heavy metals and pesticides, which are common agricultural residues. The chromoprotein allows for a convenient visible detection of these residues on agricultural produce. The chromoprotein used in all metal detection tests was mRFP which shows a visible red color. The chromoprotein used in all pesticide detection tests was AmilCP which shows a visible blue color.

In the future, we would like to do further work on our prototype so that consumers may see a fast and clear color change on the produce. We want to work on amplifying the color signal in our product so there is a clearer color change. We could experiment with different liquid viscosities and observe which consistency can best stick onto the produce. Then, we would test the functionality of our proteins in the various liquids. Moreover, we would like to come up with a new testing method for purified protein instead of using lysed cells for our experiments so it is more consumer-friendly.

## Competing Interests

The authors have declared that no competing interests exist.

## Ethics Statement

N/A

## Data Availability

Yes – all data are fully available without restriction. Sequences for the plasmids used in this study are available through the Registry of Standard Biological Parts. Links to raw data are included in Supplementary Information.

## Notes

### Competing Interest Statement

The authors have declared no competing interest.

